# Transforming the Language of Life: Transformer Neural Networks for Protein Prediction Tasks

**DOI:** 10.1101/2020.06.15.153643

**Authors:** Ananthan Nambiar, Simon Liu, Mark Hopkins, Maeve Heflin, Sergei Maslov, Anna Ritz

## Abstract

The scientific community is rapidly generating protein sequence information, but only a fraction of these proteins can be experimentally characterized. While promising deep learning approaches for protein prediction tasks have emerged, they have computational limitations or are designed to solve a specific task. We present a Transformer neural network that pre-trains task-agnostic sequence representations. This model is fine-tuned to solve two different protein prediction tasks: protein family classification and protein interaction prediction. Our method is comparable to existing state-of-the art approaches for protein family classification, while being much more general than other architectures. Further, our method outperforms all other approaches for protein interaction prediction. These results offer a promising framework for fine-tuning the pre-trained sequence representations for other protein prediction tasks.

## 1 Introduction

The advent of new protein sequencing technologies has accelerated the rate of protein discovery [1]. While protein sequence repositories are growing exponentially, existing methods for experimental characterization are not able to keep up with the present rate of novel sequence discovery [2, 3]. Currently, less than 1% of all amino acid sequences in the UniProtKB database have been experimentally characterized [2]. The explosion of uncharacterized proteins presents opportunities in computational approaches for protein characterization. Harnessing protein sequence data to identify functional characteristics is critical to understanding cellular functions as well as developing potential therapeutic applications [4]. Sequence-based methods to computationally infer protein characteristics have been critical for inferring protein function and other characteristics [5]. Thus, the development of computational methods to infer protein characteristics (which we generally describe as “protein prediction tasks”) has become paramount in the field of bioinformatics and computational biology. Here, we develop a Transformer neural network to establish task-agnostic representations of protein sequences, and use the Transformer network to solve two protein prediction tasks.

### 1.1 Background: Deep Learning

Deep learning, a class of machine learning based on the use of artificial neural networks, has recently transformed the field of computational biology and medicine through its application towards long-standing problems such as image analysis, gene expression modeling, sequence variant calling, and putative drug discovery [6, 7, 8, 9, 10]. By leveraging deep learning, field specialists have been able to efficiently design and train models without the extensive feature engineering required by previous methods. In applying deep learning to sequence-based protein characterization tasks, we first consider the field of natural language processing (NLP), which aims to analyze human language through computational techniques [11]. Deep learning has recently proven to be a critical tool for NLP, achieving state-of-the-art performance on benchmarks for named entity recognition, sentiment analysis, question answering, and text summarization, among others [12, 13].

Neural networks are functions that map one vector space to another. Thus, in order to use them for NLP tasks, we first need to represent words as real-valued vectors. Often referred to as *word embeddings*, these vector representations are typically “pre-trained” on an auxiliary task for which we have (or can automatically generate) a large amount of training data. The goal of this pre-training is to learn generically useful representations that encode deep semantic and syntactic information [12]. Then, these “smart” representations can be used to train systems for NLP tasks for which we have only a moderate amount of training data.

Word embeddings can be broadly categorized as either context-free or contextualized. Methods such as Skip-Gram (as popularized by the seminal software package word2vec) and GloVe generate context-free word vector representations [14, 15]. Once trained, these methods assign the same embedding to a given word independent of its context. Contextualized embedding models such as OpenAI-GPT [16] and Embedding from Language Model (ELMo) [17] were later developed to generate on-demand contextualized embeddings given a word and its surrounding sequence. Contextualized word representations have been shown to have superior performance to context-free representations on a variety of benchmarks [17], and have become the new standard practice in NLP.

Building upon these methods, Google recently developed a state-of-the-art contextualized technique called Bidirectional Encoder Representations from Transformers, or BERT [18], to create “deeply bidirectional” contextualized language embeddings. BERT’s architecture stacks multiple Transformer encoder layers [19], whose architecture we explain in Section 2.2, to train a single bidirectional model that generates token representations by simultaneously incorporating the leftward and rightward context.

### 1.2 Applying Deep Learning to Protein Prediction Tasks

Because of the unprecedented success in applying deep learning to NLP tasks, one of the recent interest areas in computational biology has been applying NLP-inspired techniques to amino acid sequence characterization. These techniques typically treat amino acids as analogous to individual characters in a language alphabet. Following the pre-training regime established in NLP, tools such as seq2vec and ProtVec have been developed to create amino acid sequence embeddings for protein prediction tasks. These two representations methods are based on the ELMo model and Skip-Gram technique, respectively, and demonstrate state-of-the-art accuracy when applied to bioinformatics tasks such as protein family classification, structure prediction, and protein-protein interaction prediction [20, 21].

Because of its ability to achieve higher new state-of-the-art results on a variety of NLP tasks through benchmarks such as the General Language Understanding Evalution (GLUE) benchmark, the Stanford Question Answering Dataset (SQuAD v2.0), and the Situations With Adversarial Generations (SWAG) dataset, BERT has made a breakthrough as a deeply bidirectional model [18]. Given this demonstrated success in NLP, we pre-train a similar Transformer network on amino acid sequence representations for protein prediction tasks and compare its performance to current state-of-the-art methods. Instead of using the original BERT training procedure, we use a more recent procedure called A Robustly Optimized BERT Pretraining Approach (RoBERTa) [22]. The main differences between BERT and RoBERTa lie in the pre-training step, which we highlight later in Section 2.3. We evaluate the performance of our sequence representations on two protein prediction tasks, as described below: (1) protein family classification and (2) binary protein-protein interaction prediction.

#### 1.2.1 Task: Protein Family Classification

Protein families are groups of evolutionarily-related proteins that typically share similar sequence, structural, and functional characteristics. By taking advantage of the evolutionary conservation of amino acid sequence and structure, resources such as CATH-Gene3D, PANTHER, Pfam, and SUPERFAMILY cluster amino acid sequences that share an inferred origin into hierarchical groups known as protein superfamilies, families, and subfamilies [23, 24, 25, 26]. Traditionally, classification has required the comparison of experimentally identified characteristics. However, classification methods have also been developed to computationally classify proteins based solely on sequence similarity. This approach enables us to infer functional and structural characteristics of proteins in a high-throughput manner.

Current computational approaches for protein family classification include methods such as BLASTp and profile hidden Markov models (pHMMs) which compare sequences to a large database of pre-annotated sequences. However, inference using these alignment-based methods is computationally inefficient, as they require repeated comparison of sequences to an exponentially growing database of labeled family profiles and are limited by expensive, manually-tuned processing pipelines [27]. With the exponential growth of protein discovery, the development of more scalable approaches is required to overcome traditional bottlenecks [28].

Guided by a deep learning framework, recent work using models based on convolutional neural network (CNN) and recurrent neural network (RNN) architectures has shown success in achieving state-of-the-art accuracy on the protein family classification task [28, 21]. However, these methods still produce task-specific models that are unable to generalize towards a broader range of protein prediction tasks. A recent model used an RNN architecture in a similar pre-training and fine-tuning framework to predict whether a given protein is contained in the same superfamily or fold of a reference protein [29]. However, such an approach requires pairwise comparison of sequences against multiple reference proteins, which may not be entirely representative of a protein family. Instead, we present two variants for the protein family classification task: (1) a multi-class classification problem of predicting a family label for a given sequence and (2) a binary classification problem of predicting whether sequences are members of a chosen family label.

#### 1.2.2 Task: Protein-Protein Interaction (PPI) Prediction

The next protein prediction task we highlight is protein-protein interaction (PPI) prediction. Defined as physical contacts involving molecular docking between proteins in a specific context, PPIs are fundamental to most cellular processes [30]. Research has shown that proteins, which comprise the functional machinery of cells, do not act on their own in most cases. Instead, through PPIs, proteins form physical complexes which, acting as molecular machines, are responsible for processes such as cell-cell signaling, immune response, gene expression, and cellular structure formation [31].

Identifying protein-protein interactions and creating PPI networks, or interactome networks, has thus become central to the study of biological systems [30]. By mapping these relationships, we can understand the complex interactions between individual components in a living system with a holistic approach. For instance, through the comparative analysis of PPIs in both healthy and diseased states, we can study disease mechanisms as a whole and identify potential therapeutic targets [32, 33]. However, experimental identification of PPIs has proven to be a complex and time-consuming process, thus creating the need for an efficient and reliable method of computationally predicting PPIs.

By curating data from experimental interaction studies, many of which use high-throughput techniques, public PPI resources such as APID, BioGRID, IntAct, mentha, and MINT enable the study and mapping of known binary interactions across multiple organisms. These resources detail characteristics such as interaction type, confidence scores, detection method, common pathways, and binding affinities [34, 35, 36, 37, 38]. In addition to experimentally observed PPIs, resources such as STRING collate inferred PPIs using specialized prediction pipelines [39].

We define PPI prediction as a binary classification task to predict whether two proteins will interact given their amino acid sequences. Traditionally, computational identification of PPIs has relied on genomic, structural, or domain information of the interacting proteins [40]. However, such knowledge is not readily available for most proteins. Instead, sequence-based identification currently relies on domain-based methods such as support vector machines and random forest classifiers which extract features such as amino acid distributions and domain compositions. These current approaches have limited information extraction capability and demonstrate low prediction accuracy [34, 41, 42].

More recent work using deep learning involves models that leverage architectures such as stacked autoencoders, recurrent neural networks (RNNs), and recurrent convolutional neural networks (RCNNs) [4, 43, 44]. These models have achieved state-of-the-art accuracy in the binary PPI classification task, as well as the ability to generalize to similar PPI characterization tasks such as interaction type prediction and binding affinity estimation [4, 43]. Despite this success, they still demonstrate an inability to transfer learned knowledge to more general protein prediction tasks.

### 1.3 Contributions

We propose using a Transformer neural network, which we call Protein RoBERTa (PRoBERTa) after the RoBERTa training procedure, to pre-train task-agnostic vector representations of amino acid sequences. We then fine-tune these representations towards two protein prediction tasks: *protein family classification* and *binary PPI prediction*. We compare PRoBERTa to current state-of-the-art methods for both tasks.

In the protein family classification task, we apply our model to both the multi-class classification problem and the binary classification problem. We show that the embeddings produced by PRoBERTa can be used to produce models for family classification that contain more information about protein family membership than the pre-trained embeddings, and have comparable performance to current methods that use specialized task-specific architectures.

Additionally, by evaluating PRoBERTa on the binary PPI prediction task, we demonstrate how our trained sequence embeddings can generalize to other protein prediction tasks. In the PPI prediction task, PRoBERTa outperforms other state-of-the-art methods in two classification settings. To the best of our knowledge, this is the first reported application of a Transformer network for the protein family classification and binary PPI prediction tasks.

PRoBERTa is also much more computationally efficient than recent work that applies Transformer networks to encode protein sequences to predict protein secondary structure. Using a BERT-based model, Rives et. al (2019) pre-trained their model on 128 NVIDIA V100 GPUs for 4 days [45]. In comparison, we pre-train PRoBERTa on 4 NVIDIA V100 GPUs in 18 hours using (1) a modified architecture, (2) the RoBERTa training procedure [22], and (3) the LAMB optimizer [46]. By using this framework, we are able to use a smaller pre-training corpus while obtaining state-of-the-art accuracies, increasing the computational efficiency for pre-training by a factor of 170 compared to the most recently published model.

## 2 Methods

We treat proteins as a “language” and draw ideas from the state-of-the-art techniques in natural language processing to obtain a vector representation for proteins. For a sequence of amino acids to be treated as a sentence, the alphabet of the language is defined to be the set of symbols

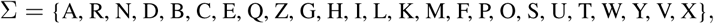

where each symbol represents one of 22 amino acids as well as three additional symbols (B, X and Z). Two of these are reserved for when it is not possible to differentiate between asparagine/aspartic acid (B) and glutamine/glutamic acid (Z). The last symbol (X) is used for unknown amino acids. This convention is based on the official IUPAC amino acid one-letter notation [47].

### 2.1 Tokenization with Byte Pair Encoding

Before amino acid sequences can be interpreted as a language, we must first define what a word is. This is more challenging for proteins than most natural languages because unlike the space character in languages like English, there is no single character (or amino acid) that is used to divide parts of an amino acid sequence into meaningful chunks. In the past, deep learning models have either used individual amino acids as input [4, 48] or have chosen to group every three amino acids as a “word” [20]. However, there has been recent interest [49] in statistically determining segments of amino acids to be used as inputs for downstream machine learning algorithms using an NLP method called byte pair encoding (BPE) [50]. Byte pair encoding was originally developed as a compression algorithm although it has been adapted more recently as a method for identifying subword units [51].

In our application, given an amino acid sequence *s* = ⟨*σ*_1_, …, *σ*_*n*_⟩ such that *σ*_*i*_ ∈ Σ, a *tokenization function* is a mapping *τ* such that *τ* (*s*) = ⟨*t*_1_, *t*_2_, …, *t*_*n*_⟩ and each *t*_*i*_ is a nonempty subsequence of *s* such that *s* = *t*_1_ … *t*_*n*_ The BPE algorithm iteratively looks for the most frequent pair of tokens and merges them to form a new token [50, 52].

### 2.2 Transformer Network Architecture

PRoBERTa uses the language representation model architecture called Bidirectional Encoder Representations from Transformers (BERT) [18]. Inspired by the BERT architecture, PRoBERTa consists of (1) an embedding layer, followed by (2) *T* = 5 stacked Transformer encoder layers, and (3) a final layer which constructs a task-specific output (Figure 1). By stacking multiple Transformer encoder layers, the aim is to capture complex higher-level information and relationships from the amino acid sequence. In total, our model has approximately 44M trainable parameters.

**Figure 1:**
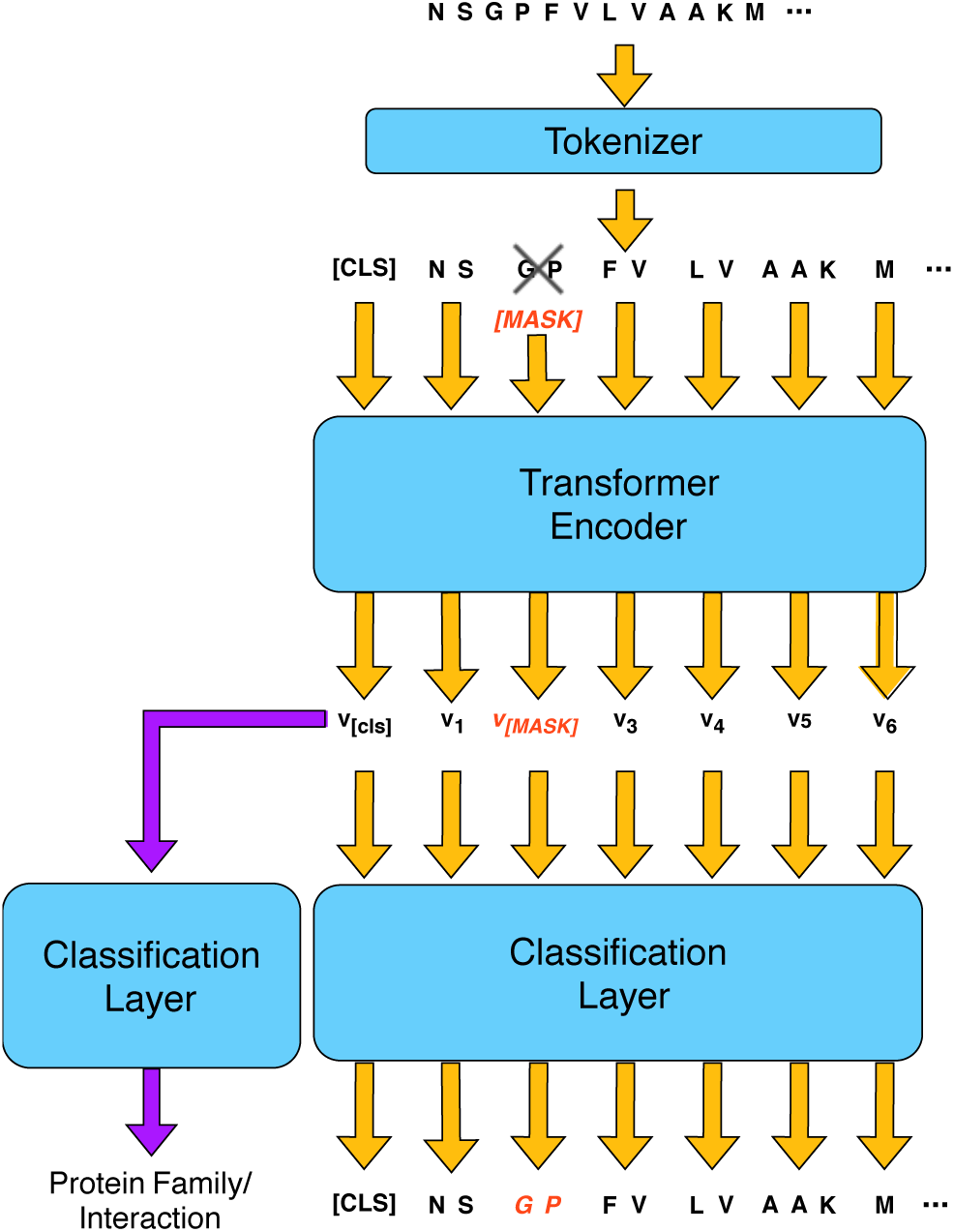
PRoBERTa pre-training and fine-tuning.

#### Model Input

A tokenized amino acid sequence ⟨*u*_1_, *u*_2_, …, *u*_*n*_⟩ is either truncated or padded to a fixed-length sequence of 512 tokens. Concretely, the model input *t* = ⟨*t*_1_, *t*_2_, …, *t*_512_⟩ is defined:

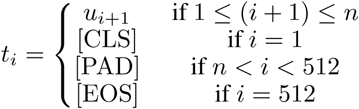

where [CLS], [PAD], and [EOS] are reserved symbols.

#### Embedding Layer

To prepare an input sequence *t* = ⟨*t*_1_, *t*_2_, …, *t*_*n*_⟩ for the Transformer encoder layers, we train an embedding layer which independently converts each token *t*_*i*_ into a vector with dimension *d* = 768. Because the model does not contain any convolution or recurrence, we incorporate sequence order information by adding positional encodings to the input embedding vectors. The specific sinusoidal function used to generate these positional encodings is chosen to allow the model to learn both the absolute as well as relative positioning of vectors [19].

#### Transformer Encoder Layer

Each Transformer encoder layer contains two sub-layers – a multi-head self-attention mechanism and a fully-connected feed-forward network – with residual connections around each sub-layer followed by a layer normalization operation [53]. Each sub-layer, and thus the entire encoder layer, takes as input and produces a list of *n* vectors, each of dimension *d* = 768.

Given an input list of vectors *x* = ⟨*x*_1_, *x*_2_, …, *x*_*n*_⟩, each vector *x*_*i*_ first travels through the multi-head self-attention mechanism. This mechanism is composed of *a* = 12 separate randomly initialized attention heads, which are trained to identify and then focus on certain subsets of positions in *x* based on their computed context relevance to *x*_*i*_. Using this mechanism, the sub-layer encodes context information from each vector *x*_*j*_ in *x*, weighted by its relevance to *x*_*i*_, into an output vector *y*_*i*_.

The initial input vector *x*_*i*_ is then added to the output vector *y*_*i*_, after which *y*_*i*_ undergoes a layer-normalization step and passes through a fully-connected feed-forward network which has a single hidden layer of size *h* = 3072 and uses a GeLU activation [54]. Each vector *y*_*i*_ passes independently through the same feed-forward network to generate the output vector *z*_*i*_. The vector *y*_*i*_ is then added to *z*_*i*_, after which *z*_*i*_ undergoes another layer-normalization step. The output for the entire Transformer layer is the list of vectors ⟨*z*_1_, *z*_2_, …, *z*_*n*_⟩ [19]

#### Model Output

Without adding any task-specific heads to the architecture, the model output is a list of *l* = 512 vectors, each with length *d* = 768. The first vector, which corresponds to the special [CLS] token, acts as an aggregate sequence representation which we use for sequence classification tasks. We refer to the entire output as the deep representation of the input amino acid sequence.

### 2.3 Model Pre-training

Following the BERT framework, we train PRoBERTa in two stages: *pre-training* and *fine-tuning* [18]. In the pre-training stage, our objective is to train the model to learn task-agnostic deep representations that capture high-level structure of amino acid sequences.

Based on the RoBERTa procedure, we pre-train PRoBERTa using only the unsupervised Masked Language Model (MLM) task, which adds a Language Modeling head to the network architecture. MLM randomly masks certain tokens and then trains the network to predict their original value [22]. We expect MLM to be a useful task because it will train the neural network to predict groups of amino acids in a sequence based on the other amino acids and their order in the sequence. This should impart some general-purpose, biologically relevant information regarding the sequences to the network. Specific training hyperparameters and optimization are detailed in Section 2.5.

Given a tokenized input sequence ⟨*t*_1_, *t*_2_, …, *t*_*n*_⟩, we select a random sample of tokens in the sequence to be masked by replacing the tokens with a special token [MASK]. Then we use cross-entropy loss to train the network to predict the masked tokens. Unlike the original BERT procedure, following the RoBERTa procedure, we generate a new masking pattern every time we feed a sequence to the model [22]. The original BERT procedure also included a Next Sentence Prediction (NSP) task. However, given that proteins are not made up of multiple sentences, as we have defined them, the NSP task is not an appropriate pre-training task. In addition, it has been shown that dropping the NSP task improves downstream performance [22].

#### Pre-training Data

We use UniProtKB/Swiss-Prot (450K unique sequences with a mean tokenized length of 129.6 tokens), a collection of experimentally annotated and reviewed amino acid sequences [2]. Sequences are tokenized with the BPE algorithm described in Section 2.1. In our experiments, the maximum vocabulary size was set to be 10,000 because we empirically observed that the average token length increased very little beyond 10,000 tokens, indicating that most of the long tokens were detected in the first 10,000.

### 2.4 Model Fine-tuning

The pre-trained model can then be leveraged as input for a wide range of downstream protein prediction tasks. In the fine-tuning stage, we first initialize the model with the trained parameters from the pre-training stage. We then modify the pre-trained architecture by replacing the original output layer with a task-specific layer with dimensions tailored to the specific task. Parameters are fine-tuned using labeled data from the prediction tasks.

Here, we fine-tune the pre-trained model for two specific prediction tasks: family classification and protein-protein interaction prediction. For our selected tasks, we feed the aggregate sequence representation corresponding to the special [CLS] token, as described in Section 2.2, into an output layer, which consists of a single-layer feed-forward neural network and softmax classifier. However, this added architecture can be customized for a wide range of protein prediction tasks other than the two we demonstrate.

#### 2.4.1 Task: Protein Family Classification

For this task, we perform two modes of classification: *binary family classification* and *multi-class family classification*. In binary family classification, we train a separate classifier for each protein family to identify which sequences belongs to a given family. This classifier performs logistic regression on the trained sequence representations from the pre-trained model. We create a balanced training dataset for each classifier consisting of all the positive samples and the same number of negative samples drawn uniformly at random without replacement from outside the family. In multi-class family classification, we train a single classifier that outputs a probability distribution over the set of all protein families. In both classification modes, we require amino acid sequences to have membership in only one protein family for ease of classification.

##### Fine-tuning Data

For the Family Classification tasks, we use 313,214 unique protein sequences from UniProtKB/Swiss-Prot whose manually curated annotations include protein family information [2].

#### 2.4.2 Task: PPI Prediction

Given a pair of tokenized amino acid sequences

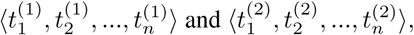

we pack them together into a single input sequence separated by a special token [SEP], which in the RoBERTa procedure is composed of two [EOS] tokens. The input representation becomes

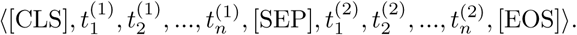

We truncate each tokenized amino acid sequence to 254 tokens before concatenation so the maximum combined length of the input sequence after the addition of the special tokens is *l* = 512 tokens.

For this problem, we use the fine-tuning procedure to add a binary classifier layer to the existing pre-training architecture. Similar to previous prediction frameworks, we treat PPI prediction as a binary classification problem. [4].

##### Fine-tuning Data

For the PPI Prediction task, we use experimentally determined human PPIs from the HIPPIE database that are confidence scored and functionally annotated [55]. Because HIPPIE only contains positive examples for PPIs, we generated two sets of negative interactions. In the “conservative” scenario, we generated 411,428 negative examples using randomly-selected pairs of human proteins from UniProt [56], resulting in a PPI dataset of 600,563 positive and negative interactions. In the “aggressive” scenario, we generated 275,402 negative examples using randomly selected pairs of proteins from HIPPIE that are not connected in the network, resulting in a PPI dataset of 536,545 positive and negative interactions. This random generation of negative examples is made possible because of the assumption that protein-protein interaction networks are sparse, though we note that there is a small chance that negatives in the aggressive scenario may be valid, but overlooked, interactions.

### 2.5 Hyperparameters and Optimization

Our two models use the same hyperparameters and optimization values, with differences between the pre-training and fine-tuning stages described below. We use the fairseq toolkit to train and evaluate our model [57]. We train our model with the LAMB optimizer [46], which is a layerwise adaptive large batch optimization technique developed for attention-based models to increase performance and reduce training time. We use the hyperparameter values from the original LAMB implementation: *β*_1_ = 0.9, *β*_2_ = 0.999, *ϵ* = 1e-8, and weight decay rate *λ* = 0.01.

We use a minibatch size of 8192 sequences with a maximum length of 512 tokens. For the pre-training stage, the learning rate is warmed up linearly over the first 3,125 updates to the peak value of 0.0025. For the fine-tuning stage, the learning rate is warmed up over the first 312 updates. Afterwards, it is adjusted using a polynomial decay policy. We take these hyperparameter values from the original LAMB implementation [46].

Per the original RoBERTa training procedure, we use a dropout of 0.1 on all Transformer layers and attention weights and a GELU activation function [18]. To avoid overfitting and balance model performance with computational efficiency, we use early stopping with a patience value of 3 (training stops after 3 consecutive epochs with no improvement in either Masked Language Model validation loss during the pre-training stage or task-specific validation accuracy during the fine-tuning stage).

### 2.6 Evaluation Metrics

For both protein prediction tasks, we use the following metrics to evaluate a model’s prediction.

1. **Accuracy** is the proportion of predictions made by the model that are correct.
2. **Precision** is the proportion of positive predictions made by the model that are correct. A high precision value indicates that the model produces few false positives.
3. **Recall** is the proportion of correct positives that are identified by the model. A high recall value indicates that the model produces few false negatives.
4. **Normalized Mutual Information (NMI) score** measures the similarity between two sets of labels for the same data. For two labels *U, V*, the NMI score is computed as

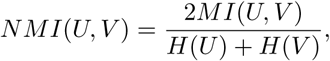

where *H*(*U*) is the entropy of *U* and *MI*(*U, V*) is the mutual information between *U* and *V*. The NMI score ranges from 0.0 (no mutual information) to 1.0 (perfect correlation), and it does not depend on the absolute values of the labels [58].

## 3 Results

We first describe the sequence features learned from the pre-trained model, before the fine-tuning stage. We then show PRoBERTa’s performance when the model is fine-tuned for the Protein Family Classification and PPI Prediction tasks. Finally, we perform a robustness analysis by limiting the amount of labeled input data during fine-tuning.

### 3.1 Protein Embeddings from the Pre-Trained Model

We pre-trained the PRoBERTa model as described in Section 2.3 on 4 NVIDIA V100 GPUs in 18 hours. We first asked whether the pre-trained model contained any biological meaning in the amino acid sequences. We created protein embeddings by concatenating the vectors of each protein’s first 128 tokens, and plotted the first two principal components of thirty proteins from seven different but related protein families (Figure 2a). We concatenated the vectors because the concatenated vectors appeared to provide better visualization results than the [CLS] token. The pre-trained model is already able to distinguish between these protein families.

**Figure 2:**
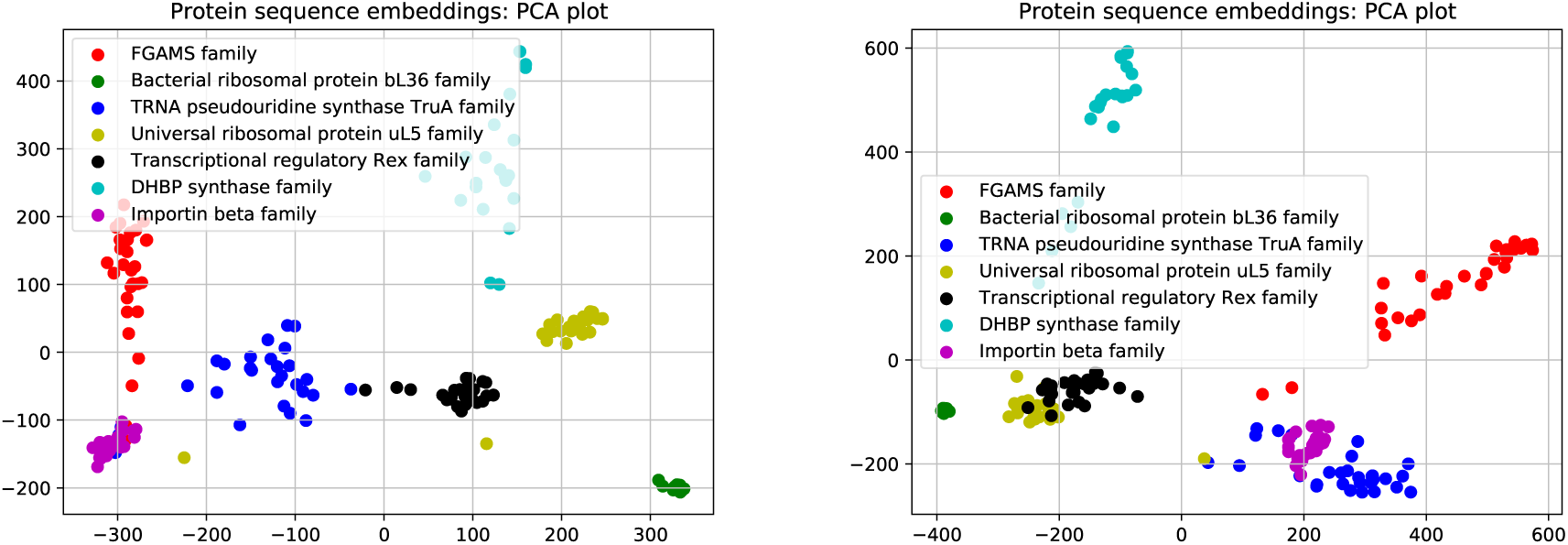
(a) The first two principal components of pre-trained embeddings for 189 amino acid sequences and (b) the first two principal components of embeddings fine-tuned on protein family classification for the same proteins

To systematically evaluate how well the pre-trained protein embeddings distinguish protein families, we clustered 9,151 protein embeddings from the manually-annotated human proteins in UniProt and compared the clusters to the 3,860 annotated protein families. However, because this unsupervised learning approach is especially susceptible to the curse of dimensionality, we substituted the model described in Section 2.2 with a similar model but with embedding dimension *d* = 192 instead of *d* = 768 [59] and summed the vectors for each token in the amino acid sequence. This lower dimensional embedding is only used for the unsupervised learning task. To cluster, we reduced the pre-trained protein embeddings to twenty dimensions using PCA and applied *k*-means clustering using Euclidean distance and setting *k* = 4000 to approximately reflect the number of annotated families. The clusters largely correlate to the protein families, as shown by five randomly-selected clusters from the 200 largest clusters (Figure 3). The mean NMI of the pre-trained vectors is 0.798213 averaged over 20 runs, which is significantly larger than than the NMI of randomly assigned clusters (Figure 4, *t*-test *p*-value: 8 × 10^−180^).

**Figure 3:**
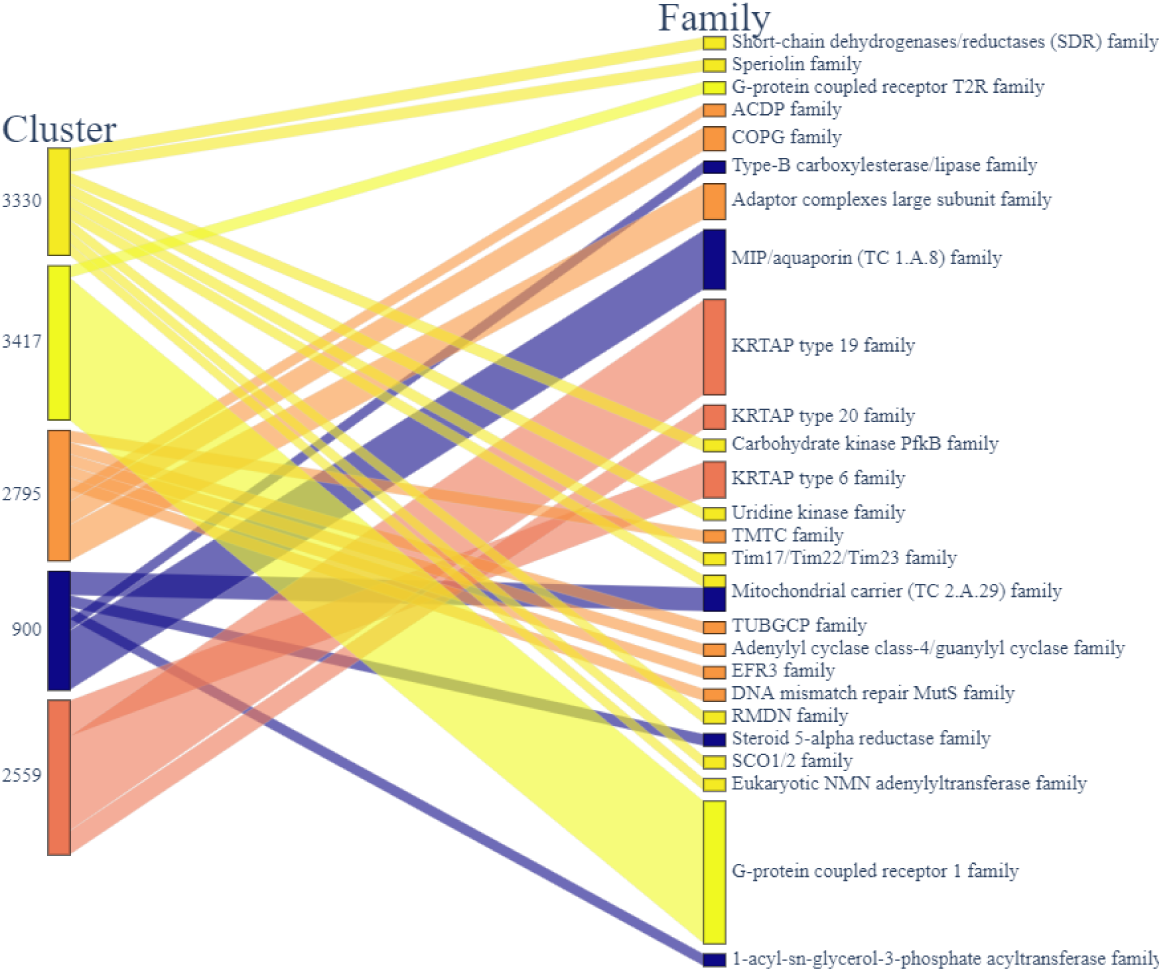
Parallel set plot of 5 clusters randomly selected from the 200 largest clusters.

**Figure 4:**
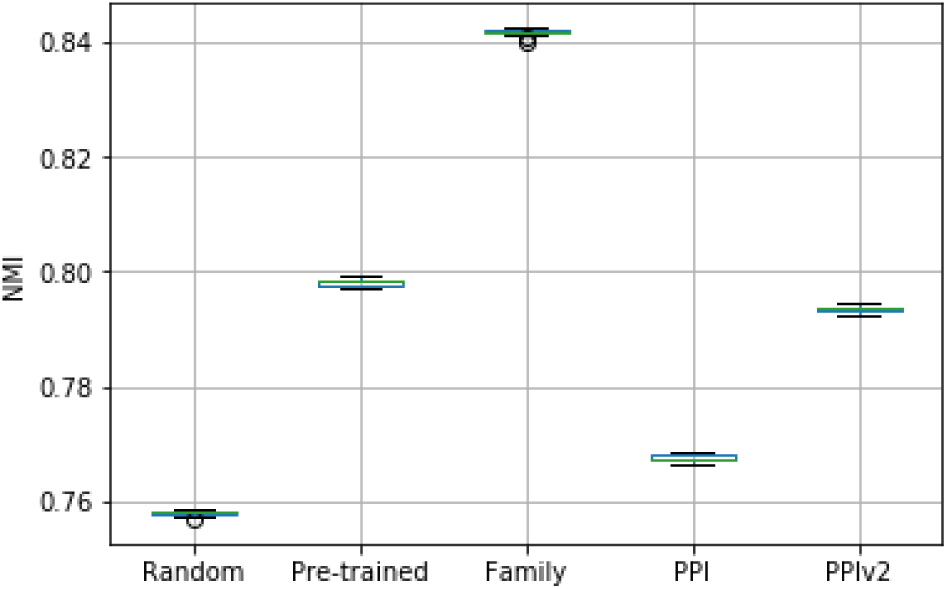
Plot of NMI values comparing the unsupervised clustering on three different versions of PRoBERTa embeddings with the true protein families given by UniProt. PPI: conservative scenario; PPIv2: aggressive scenario.

### 3.2 Protein Family Classification

Given the promise of the protein embeddings, we then evaluated how well the PRoBERTa model performs on the protein family classification task (Section 2.4.1). Clustering the vectors fine-tuned on the protein family classification task increases the NMI even more than the pre-trained model (Figure 4), suggesting that the fine-tuned embeddings have more specific information related to protein classification.

In the binary classification task, we trained a separate logistic regression classifier for each protein family with greater than 50 proteins and measured the weighted mean accuracy as 0.98 with the lowest scoring family being made up of 57 proteins and having an accuracy of 0.77. In performing this, we randomly withheld 30% of the proteins from each family to be used as the test set. We compared PRoBERTa+logistic to three other NLP based embedding methods: ProtVec, which is a protein embedding method inspired by Word2Vec; ProtDocVec, which modifies ProtVec to use Doc2Vec; and ProtFreqVec, which simply uses the frequency of triplets of amino acids form embedding vectors [20, 60].

PRoBERTa+logistic performs better than ProtVec+logistic and has similar performance to ProtDocVec+logistic and ProtFreqVec+logistic (Table 1). In Figure 2 (a), all the families but the importin beta family were chosen from families that have accuracies that are representative of the overall accuracy (ranging from 0.96 to 1.0) while the importin beta protein family has a substantially lower accuracy of 0.77. The PCA plots in Figure 2 show that the proteins in this less accurately classified family are co-located with proteins from FGAMS family and TRNA pseudouridine synthase TruA family.

**Table 1:**
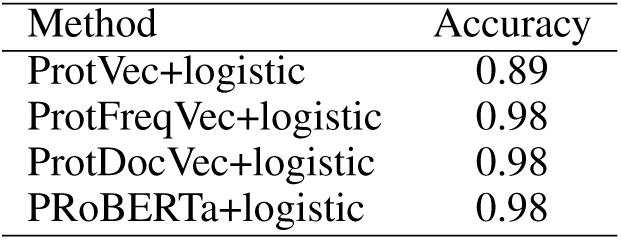
Comparison of binary family classification.

In the multi-class family classification task, we used fine-tuning to add an output layer that maps to protein family labels. This was done using the dataset of 313,214 UniProt proteins with only one associated family. These proteins were split into train/validation/test sets (0.8/0.1/0.1) and provided us with a classifier that had an accuracy of 0.92 on the test set. We then compared this to two other multi-class family classifiers including a simple CNN, made up of four convolution layers, as a base line and DeepFam, a CNN method that is the current state-of-the-art method for protein family classification. In particular, DeepFam is made up of a convolution layer with eight different kernel sizes and 250 convolution units for each kernel size. The max-pooled features extracted through these convolution units are passed through a fully-connected layer to get the predicted family [3]. PRoBERTa with a classification layer performed better than the baseline method and had comparable accuracy to DeepFam (Table 2).

**Table 2:** Comparison of multi-class family classification.

| Method | Accuracy |
| --- | --- |
| DeepFam | 0.95 |
| Simple CNN | 0.72 |
| PRoBERTa | 0.92 |

### 3.3 PPI Prediction

We next assessed PRoBERTa on the PPI task using the conservative and aggressive scenarios for sampling negative interactions (Section 2.4.2).

#### 3.3.1 Conservative Scenario

We first evaluated how well the fine-tuned model’s embeddings captured protein families compared to the pre-trained embeddings. In the conservative scenario, the fine-tuned model produces embeddings that cluster with a lower NMI with the protein families compared to the pre-trained embeddings (Figure 4), indicating that the parameters of the model fine-tuned on predicting interactions are not as tuned to protein family classification. We found that the fine-tuned PRoBERTa PPI classifier had an accuracy of 0.96 with a precision and recall of 0.93 and 0.94, respectively, (Table 3) as well as an ROC AUC of 0.99 (Figure 5). In these runs, we only used 20% of the available training and validation data from the train/validation/test (0.8/0.1/0.1) split. That is, only 18% (108,103) of the total data is seen by the model when training. This was done because some of the methods we compare against were not able to scale up to use 80% of the data for training and thus would not be able to make a fair comparison to the PRoBERTa model trained with the entire train set. We then compared our results to PIPR, which is one of the best PPI prediction neural networks currently available, using a similar number of interactions from our dataset [4]. PIPR uses a residual convolutional neural network (RCNN) architecture to extract both sequential information as well as local features relevant for PPI prediction. We also compare our embeddings to the ProtVec embeddings combined with a neural network with three hidden layers (which we call *DeepInteract*) that predicts PPI. Finally, we try a biologically motivated transfer learning approach by first training a DeepFam network on protein family classification and then using one of the hidden layers as the vector representations of the proteins to be used by DeepInteract. As seen in in Figure 5, PRoBERTa with a classification layer outperforms all of these methods by a large margin. Further, when using the complete dataset, the accuracy reaches 0.99 (Table 3 and Figure 7).

**Table 3:** PPI prediction results using 20% of training data (top) and using 100% of training data (bottom).

| Method | Conservative |  |  | Aggressive |  |  |
| --- | --- | --- | --- | --- | --- | --- |
|  | Acc. | Prec. | Rec. | Acc. | Prec. | Rec. |
| DeepFam+DI* | 0.79 | 0.79 | 0.66 | 0.75 | 0.76 | 0.73 |
| PIPR | 0.81 | 0.75 | 0.77 | 0.77 | 0.77 | **0.77** |
| ProtVec+DI* | 0.80 | 0.78 | 0.70 | 0.73 | 0.72 | 0.76 |
| PRoBERTa | **0.96** | **0.93** | **0.94** | **0.79** | **0.82** | 0.73 |
| PRoBERTa<br>(100% training) | **0.99** | **0.97** | **0.98** | **0.84** | **0.83** | **0.84** |

\*DI: Deep Interact

**Figure 5:**
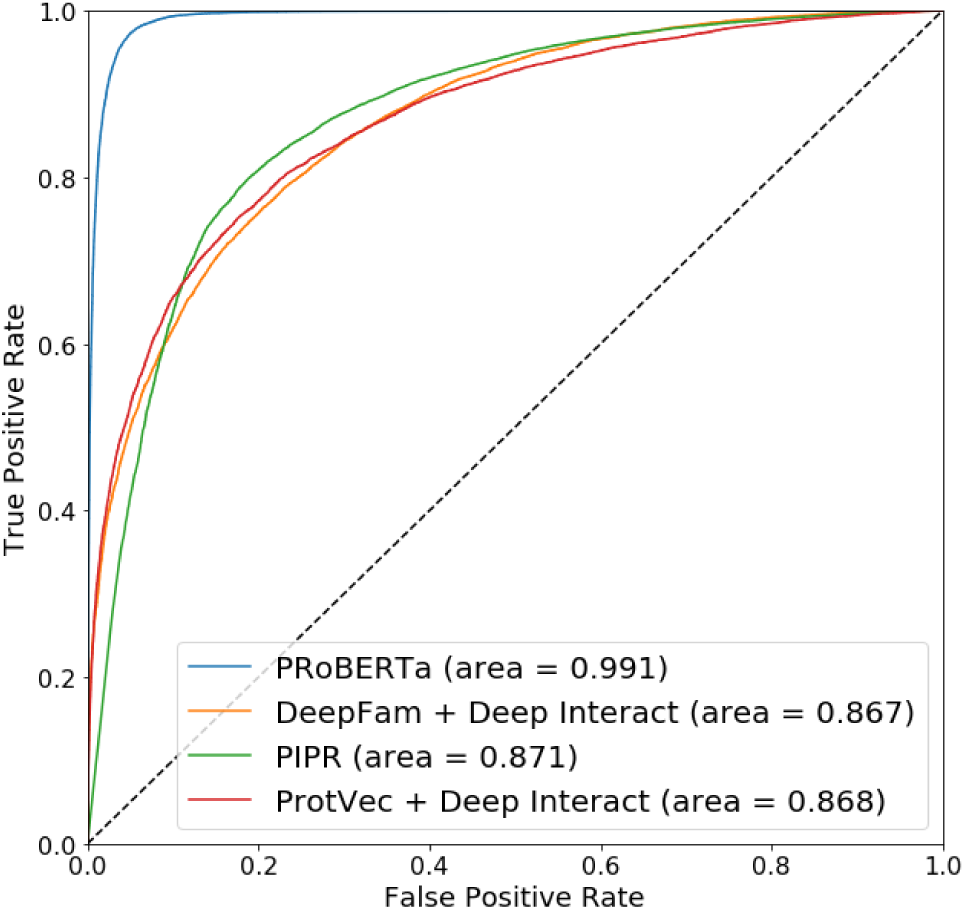
Receiver operating characteristic (ROC) curves for PPI prediction for the conservative scenario

**Figure 6:**
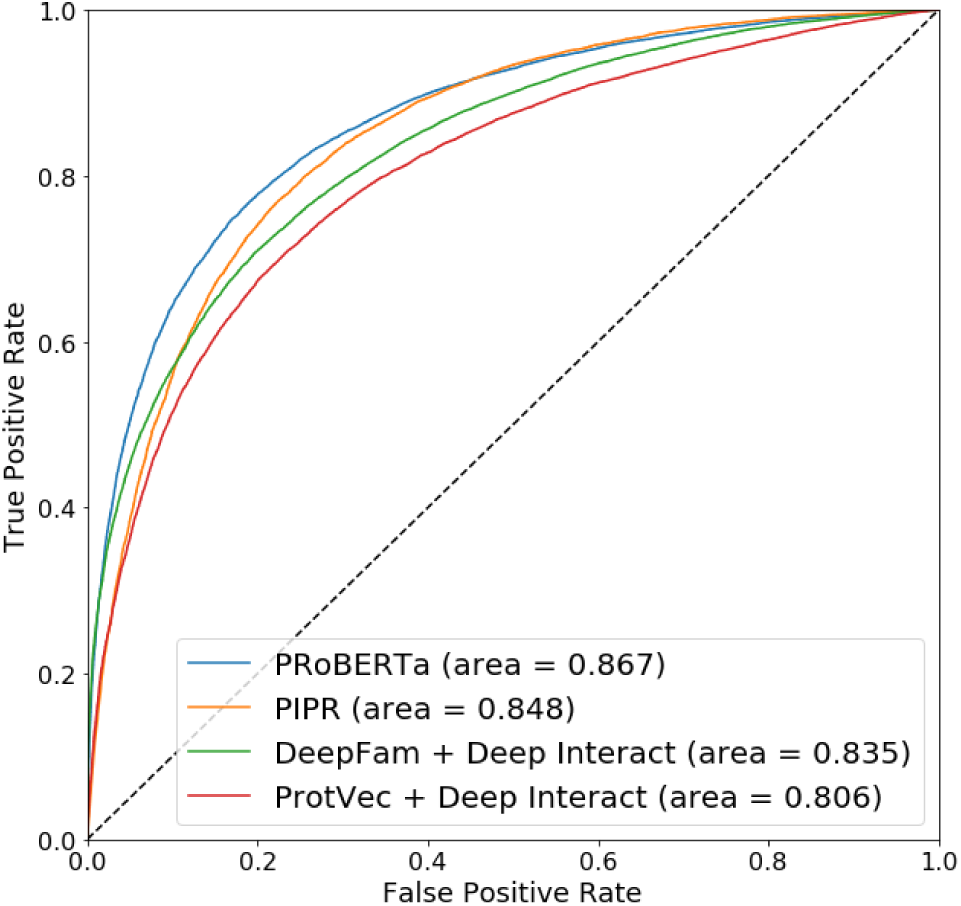
Receiver operating characteristic (ROC) curves for PPI prediction for the aggressive scenario

**Figure 7:**
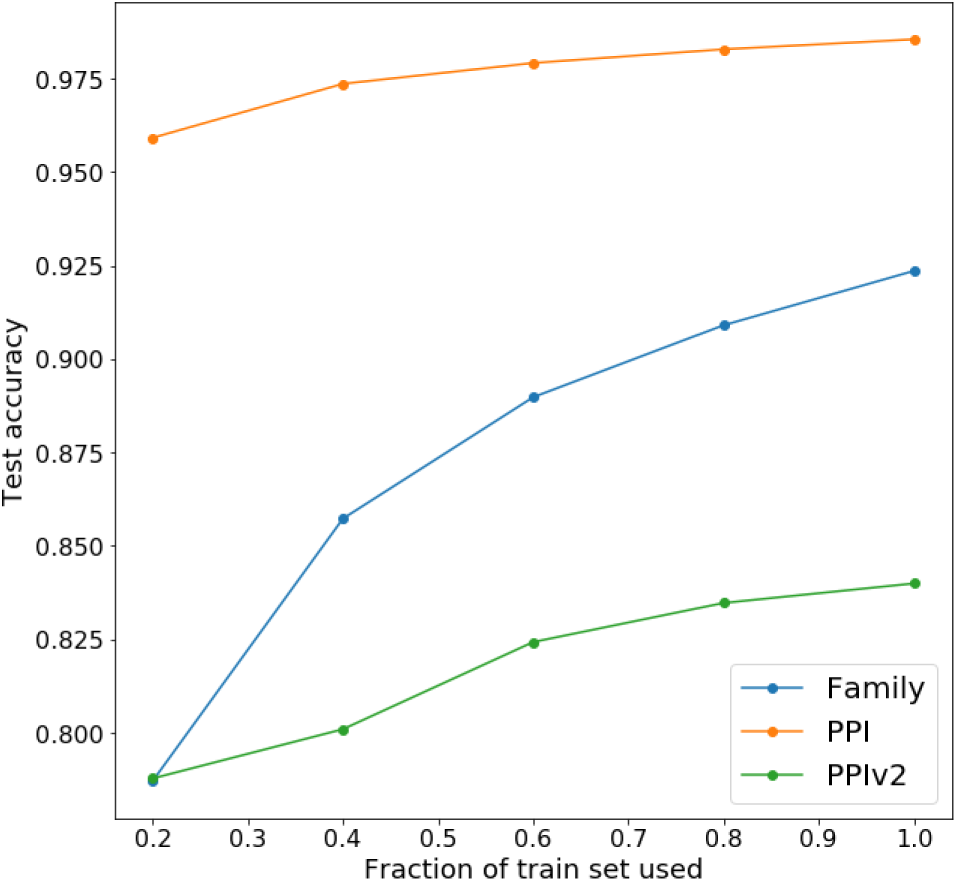
Varying amount of training data for fine-tuning. PPI: conservative scenario; PPIv2: aggressive scenario.

#### 3.3.2 Aggressive Scenario

Similar to the conservative scenario, the model fine-tuned on the aggressive PPI dataset produces embeddings with a lower NMI with protein families than the pre-trained embeddings (Figure 4). For this scenario, PRoBERTa had an accuracy of 0.79 with a precision and recall of 0.82 and 0.73, respectively, (Table 3) as well as an ROC AUC of 0.87 (Figure 6). Similar to above, we only use 20% of available training and validation data. As seen in Figure 6, PRoBERTa with a classification layer still does better than PIPR, which does better than the other two methods. Moreover, when the full training dataset is used, the accuracy of the model improves to 0.84 (Table 3 and Figure 7). However, PRoBERTa does decrease in performance and over-fits (train accuracy: 0.98, test accuracy: 0.77) for much smaller datasets such as the yeast interactions dataset for which we had only 11,164 usable interactions [4].

### 3.4 PRoBERTa Scalability and Robustness

We also investigated the robustness of the models by varying the amount of training data for both fine-tuning tasks. For the PPI prediction task in Section 3.3, we used 20% of the training data in order to compare to existing methods; here, we are able to use all of the training data, improving the accuracy for both the conservative and aggressive scenarios (Table 3).

To assess the robustness of PRoBERTa on the protein prediction tasks, we used fractions of the 0.8 split that made up the fine-tuning train set for task training. For example, if 90% of the train set was used, this meant that 90% × 0.8 or 72% of the entire dataset was used for training. Figure 7 shows the change in accuracy with different fractions of the train set used. This shows that all three models were somewhat robust to different amounts of training data. The PPI models appears to be more robust (they have smaller slopes) than the Protein Family model. However, it should be noted that the complete train set for the Protein Family model contained 250,504 proteins, while the PPI model had 480,455 interactions in the conservative scenario and 429,239 interactions in the aggressive scenario. This difference in robustness could be due to the absolute difference in number of training data points.

## 4 Discussion

In this paper, we propose a Transformer based neural network architecture, called PRoBERTa, for protein characterization tasks. We found that the pre-trained embeddings contain general yet biologically relevant information regarding the proteins and fine-tuning pushes the embeddings to have more specific information at the cost of generality.

Moreover, we found that using the embeddings for Protein Family Classification produced results that were comparable to the current best methods. In particular, we performed two different forms of classification; a binary classification that classified a protein as “in the family” or “not in the family” and a multi-class family classification. The multi-class family classification was based on the assumption that there there is only one class per protein. Proteins that belong to more than one family were excluded from this classifier but not the binary classifiers.

Furthermore, we used embeddings from PRoBERTa for a fundamentally different problem, PPI Prediction, using two different datasets generated from the HIPPIE database and found that with sufficient data, it substantially outperforms the current state-of-the-art method in the conservative scenario and still performs better than the other methods in the aggressive scenario. When evaluated on the aggressive dataset, the model trained on the conservative dataset scores an overall accuracy of 0.59, with a precision and recall of 0.54 and 0.94, respectively. This, combined with the larger decrease in NMI with protein families in the aggressive scenario (Figure 4), suggests that the model in the conservative scenario performs something closer to a protein classification task to identify which proteins are present in the HIPPIE dataset and are thus more likely to correspond to positive interaction examples.

The efficiency of PRoBERTa over existing methods (a speedup in pre-training time by a factor of 170 compared to the similar BERT-based model by Rives et. al (2019) [45]) provides unprecedented opportunities for using the growing amount of sequence data in protein prediction tasks. Further, PRoBERTa’s success in these two different protein prediction tasks alludes to the generality of the embeddings and their potential to be used in other tasks such as predicting protein binding affinity, protein interaction types and identifying proteins associated with particular diseases. In light of the COVID-19 pandemic, we are currently working on adapting PRoBERTa for vaccine design.

## 5 Acknowledgements

This work has been supported by the National Science Foundation (awards #1750981 and #1725729). This work has also been partially supported by the Google Cloud Platform research credits program (to AR, MH, and AN). AN would like to thank Mark Bedau, Norman Packard and the Reed College Artificial Life Lab for insightful discussions and Desiree Odgers for inspiring the idea of taking a linguistic approach to a biological problem.

